# Gabapentin’s Effect on Human Dorsal Root Ganglia: Donor-Specific Electrophysiological and Transcriptomic Profiles

**DOI:** 10.1101/2024.12.05.627067

**Authors:** Jenna B. Demeter, Nesia A. Zurek, Maddy R. Koch, Aleyah E. Goins, Cristian O. Holguin, Mark W. Shilling, Reza Ehsanian, Sascha R.A. Alles, June Bryan I. de la Peña

## Abstract

Neuropathic pain affects approximately 10% of the adult population and is commonly treated with gabapentin (GBP), a repurposed anticonvulsant drug. Despite its widespread clinical use, GBP’s efficacy varies significantly among patients, highlighting the need to better understand its functional and molecular impacts on human pain-sensing neurons. In this study, we characterized the electrophysiological and transcriptomic effects of GBP on primary sensory neurons derived from the dorsal root ganglia (DRG) of ethically consented human donors. Using patch-clamp electrophysiology, we demonstrated that GBP treatment reduced neuronal excitability, with more pronounced effects in multi-firing vs. single-firing neuronal subtypes. Notably, significant donor-specific variability was observed in electrophysiological responsiveness to GBP treatment *in vitro*. RNA sequencing of DRG tissue from the GBP-responsive donor revealed differences in the transcriptome-wide expression of genes associated with ion transport, synaptic transmission, inflammation, and immune response relative to non-responsive donors. Cross-transcriptomic analyses further showed that GBP treatment counteracted these altered processes, rescuing aberrant gene expression at the pathway level and for several key genes. This study provides a comprehensive electrophysiological and transcriptomic profile of the effects of GBP on human DRG neurons. These findings enhance our understanding of GBP’s mechanistic actions on peripheral sensory neurons and could help optimize its clinical use for neuropathic pain management.

## Introduction

Neuropathic pain affects approximately 10% of the adult population and arises from damage or dysfunction within the somatosensory nervous system [2,28,44]. It manifests in conditions such as diabetic neuropathy, postherpetic neuralgia, and chemotherapy-induced peripheral neuropathy (CIPN). The management of neuropathic pain poses significant challenges due to its complex pathophysiology, which involves aberrant ion channel activity, synaptic dysfunction, and neuroinflammation [3,5,23,32].

Gabapentin (GBP), originally developed as an anticonvulsant, has become one of the most widely prescribed treatments for neuropathic pain [31,48,49]. GBP exerts its effects by binding to the α2δ-1 subunit of voltage-gated calcium channels, reducing calcium influx, neurotransmitter release, and neuronal hyperexcitability [10,35,49]. Despite its widespread clinical use, GBP’s pain-relieving effects varies significantly among patients [31,48]. Additionally, side effects—including sedation, dizziness, and potential for misuse—raise concerns about its long-term utility [16,24,36,37]. These limitations underscore the importance of understanding GBP’s mechanisms of action in human nociceptors.

Despite its well-documented role in modulating calcium channels, GBP’s precise functional and molecular effects on human sensory neurons remain poorly characterized [49]. Most insights into GBP’s mechanisms derive from rodent models, which do not fully capture the complexities of human sensory neuron biology [38]. Furthermore, transcriptome-wide effects of GBP have not been systematically studied in any neuronal system, leaving a significant gap in understanding its molecular actions. Human dorsal root ganglia (hDRG) neurons, which serve as critical mediators of nociception and are damaged in peripheral neuropathies, offer a unique and clinically relevant model for studying GBP’s mechanisms of action [53].

To bridge this knowledge gap, we examined the electrophysiological and transcriptomic effects of GBP on primary hDRG. Samples were obtained from ethically consented organ donors, and GBP’s effects were evaluated using patch-clamp electrophysiology and RNA sequencing (RNA-seq). Our experiments revealed that *in vitro* GBP treatment reduces neuronal excitability, with effects more pronounced in multi-firing compared to single-firing neuronal subtypes. Transcriptomic analysis further revealed that GBP treatment modulates the expression of genes involved in ion transport, synaptic transmission, inflammation, and immune response.

Importantly, we observed significant donor-specific variability in GBP responsiveness, emphasizing the influence of intrinsic biological differences. One donor (H22, the “responder”) had more excitable hDRG neurons compared to the other two donors studied (H16/H17, the “non-responders”). GBP treatment reduced the excitability of H22 neurons to levels comparable to or lower than the non-responders. Transcriptomic data revealed that the responder has enrichment of genes associated with ion transport and voltage-gated ion channel activity, potentially explaining its heightened excitability and responsiveness to GBP. Cross-transcriptomic analyses demonstrated that GBP treatment counteracted aberrant expression of genes related to ion transport, synaptic signaling, and inflammatory and immune responses in the responder’s DRG.

This study highlights the variability in GBP responsiveness and its potential molecular underpinnings, emphasizing the need for personalized approaches to pain management. By integrating electrophysiological and transcriptomic data, we provide a comprehensive profile of GBP’s effects on hDRG neurons. These findings not only deepen our understanding of GBP’s mechanism of action but also offer valuable insights for optimizing its clinical application in managing neuropathic pain.

## Materials and Methods

### hDRG Extraction and Culture

hDRG were acquired from ethically consented organ donors at the University of New Mexico Hospital in coordination with New Mexico Donor Services. This study was conducted in compliance with institutional ethical guidelines and approved by the University of New Mexico Health Sciences Center Human Research Review Committee (approval numbers #21-412 and #23-205). Research was conducted in accordance with the Declaration of the World Medical Association. Details of the donors are summarized in **Table 1**. hDRG neuron (hDRG-N) cultures were performed as previously described [46,53]. Briefly, hDRG tissues were mechanically and enzymatically dissociated, washed, then cultured on poly-D-lysine (PDL)-coated coverslips or wells in Neurobasal Plus medium supplemented with fetal bovine serum (FBS), B-27, GlutaMAX, and antibiotic-antimycotic (ThermoFisher). Cultures were maintained for up to 11 days *in vitro* (DIV). For treatment experiments, hDRG-N cultures were exposed to 100 µM GBP (G0318, TCI Chemicals, Portland, OR) or vehicle control (equal volume H_2_O) overnight (∼16 hrs) before downstream analyses, including electrophysiological and molecular assays.

**Table 1:** Donor demographics and characteristics.

| Donor | Sex | Age<br>(years) | Ethnicity | BMI | Toxicology | Cause of death |
| --- | --- | --- | --- | --- | --- | --- |
| H16 | Male | 51 | White | 27.2 | Opioids,<br>benzodiazepines | Cerebrovascular<br>accident/ stroke |
| H17 | Female | 23 | White/<br>Hispanic | 26.6 | Opioids (fentanyl),<br>cannabinoids | Cardiac arrest, anoxia |
| H22 | Female | 37 | White | 22.5 | Amphetamines,<br>benzodiazepines | Head trauma (motor<br>vehicle collision),<br>intracranial hemorrhage |

### Immunohistochemistry

Immediately after extraction, hDRG tissues were embedded in optimal cutting temperature (OCT) compound, sectioned at 20 µm using a cryostat. Sections were mounted onto glass slides and allowed to adhere overnight at room temperature. hDRG sections were washed 5x in 1X PBS before being incubated for 1 hr at room temperature in a blocking solution containing 0.3% Triton X-100 (A16046.AE, ThermoFisher) and 10% fish gelatin in 1X PBS (22010, Biotium). The sections were incubated at room temperature with the following primary antibodies and dilutions: chicken anti-peripherin (PA1-10012, Invitrogen, 1:1000) and rabbit anti-α2δ-1 (ACC-015, Alomone Labs, 1:1000). Slides were rinsed 5x in 1X PBS, followed by a 2-hr incubation in secondary antibodies: goat anti-chicken AF555 (AB150170, Abcam, 1:2000) and donkey anti-rabbit AF488 (A21206, Invitrogen, 1:2000). After secondary antibody incubation, the sections were washed 3x in 1X PBS then treated with Sudan Black B (J62268.09, ThermoFisher) for 30 minutes to reduce autofluorescence. Coverslips were applied using Fluoromount-G Mounting Medium with DAPI (00-4959-52, Thermo Fisher). Immunofluorescence was visualized using a Leica THUNDER Imaging System (Leica Microsystems) at 40X magnification and an Olympus IXplore Spin/Yokogawa CSU-W1 Confocal Microscope (Evident Microscopy) at 60X magnification.

### Whole-cell Patch-clamp Electrophysiology

Patch-clamp electrophysiology was performed as previously described [53]. Recordings were done at room temperature and perfused with artificial cerebrospinal fluid (aCSF) bubbled with 95% O_2_/5% CO_2_. aCSF contained 113 mM NaCl, 3 mM KCl, 25 mM NaHCO_3,_ 1 mM NaH_2_PO_4,_ 2 mM CaCl_2,_ 2 mM MgCl_2_, and 11 mM dextrose (MilliporeSigma). Patch pipettes with electrode resistance of 4-9 MΩ were pulled with Zeitz puller (Werner Zeitz, Martinsried, Germany) from borosilicate thick glass (GC150F, Sutter Instruments, Novato, CA) and with an intracellular patch pipette solution containing 120 mM K-gluconate, 11 mM KCl, 1 mM CaCl_2,_ 2 mM MgCl_2,_ 10 mM HEPES, 11 mM EGTA, and 4 mM Mg-ATP. hDRG-N were identified using an IR-2000 Infrared Monochrome Video Camera (Dage MTI, Indiana City, MI) equipped with interference contrast optics. Cell diameter was measured using the IR-Capture software (Dage MTI). Current clamp recordings were done using a MultiClamp 700B Microelectrode Amplifier (Molecular Devices, San Jose, CA). Data was acquired using an Axon Digidata 1550B Low Noise Data Acquisition System (Molecular Devices, San Jose, CA) and Axon Clampex 11 software (Molecular Devices). Bridge balance was applied for all recordings. Current clamp recording sweeps started with 25 ms.

### Electrophysiology Data Analysis

Analysis was done as previously described using the Python package pyABF (v3.12) and Easy Electrophysiology (v2.5.1) [19,53]. Cell capacitance was measured using the whole-cell compensation circuit from MultiClamp 700B (Molecular Devices). Current clamp recording sweeps started with 25 ms of the cell at resting membrane potential (RMP), followed by 500 ms current input in 10 pA increments, starting at-100 pA for the first sweep, followed by 500 ms recovery between each sweep until the neuron reached inactivation or up to 4 nA. RMP was calculated for the first sweep in the 25 ms after establishing whole cell configuration. The first-100 pA hyperpolarizing current injection was used to calculate sag ratio and input resistance (R_in_). If a neuron fired an action potential (AP) during the recovery period of any hyperpolarizing current injection, the neuron was classified as rebound-firing. Rheobase was calculated as the current injection in which a neuron fired its first AP during the depolarizing 500 ms current injection; therefore, the lowest possible value for rheobase is 10 pA. The first-spike latency (FSL) was the time after the start of current injection of the rheobase AP. Neurons that had an FSL > 100 ms were classified as delayed-firing. hDRG-N that fired more than one AP during any depolarizing current injection step were classified as multi-firing while hDRG-N that fired only one AP were classified as single-firing. Spontaneous activity was measured in current clamp mode with either no applied current (at rest) or with enough current applied to hold the membrane voltage at-45 mV; cells that fired any AP during a 30 s recording were classified as having spontaneous activity. Depolarizing spontaneous fluctuations (DSFs) were analyzed using the spontaneous activity at rest trace with FIBSI [34]. AP waveform properties were calculated using the rheobase AP using Method II in Easy Electrophysiology [39]. We did not correct for liquid junction potential. Statistics comparing all electrophysiological properties were done in GraphPad Prism (v10.0.2) using a Mann-Whitney test or a Fisher’s Exact test.

### RNA Sequencing and Quality Control

Total RNA was isolated from frozen hDRG tissue and GBP- or vehicle-treated primary hDRG cultures (n = 3 per condition) using the Monarch Total RNA Miniprep Kit (New England Biolabs) following the manufacturer’s protocol. The quantity and purity of isolated RNA were initially assessed using a NanoDrop spectrophotometer (Thermo Fisher Scientific), ensuring OD 260/280 ratio ≥ 1.8, indicating no or minimal contamination. The integrity of the RNA was further evaluated using the 2100 Bioanalyzer (Agilent) (for hDRG tissue samples) or the 5400 Fragment Analyzer System (Agilent) (for hDRG culture samples), with samples achieving an RNA Integrity Number (RIN) ≥ 5.3 deemed suitable for library construction.

Total RNA with rRNA removed or poly-A-captured mRNA underwent library construction, the former of which was directional. The libraries were sequenced on a NovaSeq X Plus Sequencing System (Illumina) using a 150 bp paired-end read strategy. This sequencing approach provided comprehensive coverage, generating approximately 12 Gb (H16/H17) or 6 GB (H22, culture) of raw data per sample, which resulted in over 20 million raw read pairs for all samples. This depth of coverage ensures robust quantification of gene expression levels and accurate detection of differentially expressed genes (DEGs). Phred scores— Q20 > 97% and Q30 > 93%—for all samples indicated that sequencing was of high quality and data was suitable for further analysis.

### Bioinformatic Analysis

Raw data underwent filtering of paired reads when either read 1) was contaminated with an adapter, 2) had > 10% uncertain nucleotides (N) (rarely), or 3) was of low quality (Q5 for > 50% of nucleotides) (very rare). Poly-G tails— which occur in two-color chemistry systems when a black G is repeatedly called after synthesis has terminated—were trimmed using BBDuk when > 20 bp. Quality was ensured by several metrics using FastQC.

High-quality reads were aligned to an Ensembl human reference genome (GRCh38, release 112) using HISAT2, and gene expression was quantified using StringTie. Differential expression analyses for genes and transcripts was conducted using edgeR. The generalized linear model (GLM) quasi-likelihood (QL) method was used to analyze differential expression between the responder (H22) and non-responder (H16 and H17) hDRG tissues. All of our sequenced donor hDRG tissues that had complete accompanying metadata were included in the count data to thoroughly determine genes/ transcripts that are uniquely differentially expressed in the responder vs. non-responder given the wider transcriptomic context of all donors. To better account for likely confounding factors, library prep/ RNA-sequencing (RNA-seq) strategy and donor age (in 10-year bins), sex, and ethnicity (white and/or Hispanic) were included in the design matrix in addition to responsiveness to GBP (the contrast being made). To analyze differential expression between GBP- and vehicle-treated hDRG cultures, the quantile-adjusted conditional maximum likelihood (qCML) method was used since treatment was the only factor. A threshold of |log2FC| ≥ 0.04 and p ≤ 0.05 was used to define significant DEGs. This Ensembl-HISAT2-StringTie-edgeR pipeline was recently ranked in an RNA-seq benchmarking study as the overall most accurate pipeline for identifying DEGs [47].

Gene Ontology (GO) gene set enrichment analysis (GSEA) was conducted using clusterProfiler for both the responder vs. non-responder and GBP- vs. vehicle-treated differential expression analyses, using log2FC-ranked lists as input. A threshold of p ≤ 0.05 was used to define significantly enriched terms. While significantly enriched terms representing a variety of biological processes, molecular functions, and cellular compartments were identified, key representative terms were selected along the themes of ion transport, synaptic transmission, inflammation, and immune response given the ubiquity of these themes in the significantly enriched term sets as well as their relevance to neuronal activity and pain modulation.

Scaled PCA analysis of hDRG tissue samples was conducted in base R using counts that underwent trimmed mean of M-values (TMM) normalization. The heatmap of strong DEGs (|log2FC| ≥ 0.4, FDR ≤ 0.1) in GBP- vs. vehicle-treated hDRG cultures was generated using pheatmap, with log2TPM as input and scaling by gene to generate Z-scores for each sample. All other results were visualized using ggplot2.

## Results

### α2δ1 is abundantly expressed in hDRG neurons

α2δ1 (encoded by *CACNA2D1*) is the canonical target of GBP, playing a critical role in its mechanism of action. To confirm expression of α2δ1 in our hDRG samples, we analyzed single-cell RNA sequencing data from recently published datasets on human DRG neurons [7]. Cells were divided into clusters based on the expression of marker genes, allowing us to classify neuronal subtypes (Figure 1A). Analysis of *CACNA2D1* expression across these clusters revealed that the receptor is ubiquitously expressed in human DRG neurons (Figure 1B). Notably, within the neuronal population, α2δ1 was broadly expressed across all neuronal subtypes, with varying levels of expression in specific subgroups: Ntrk3^high^+Ntrk2 (60%), Pvalb (72.09%), Atf3 (76.92%), Calca+Bmpr1b (84%), Trpm8 (85.71%), Ntrk3^high^+S100a16 (87.50%), Calca+Sstr2 (89.74%), Sst (93.10%), Mrgprd (96%), Calca+Smr2 (96.88%), Calca+Adra2a (100%), Calca+Oprk1 (100%), Ntrk3^low^+Ntrk2 (100%), and Th (100%) (Figure 1C). These data demonstrate that α2δ1 is not only broadly expressed but also consistently present in various nociceptive and mechanosensory neuronal subtypes, suggesting its pivotal role across sensory neuron populations.

**Figure 1:**
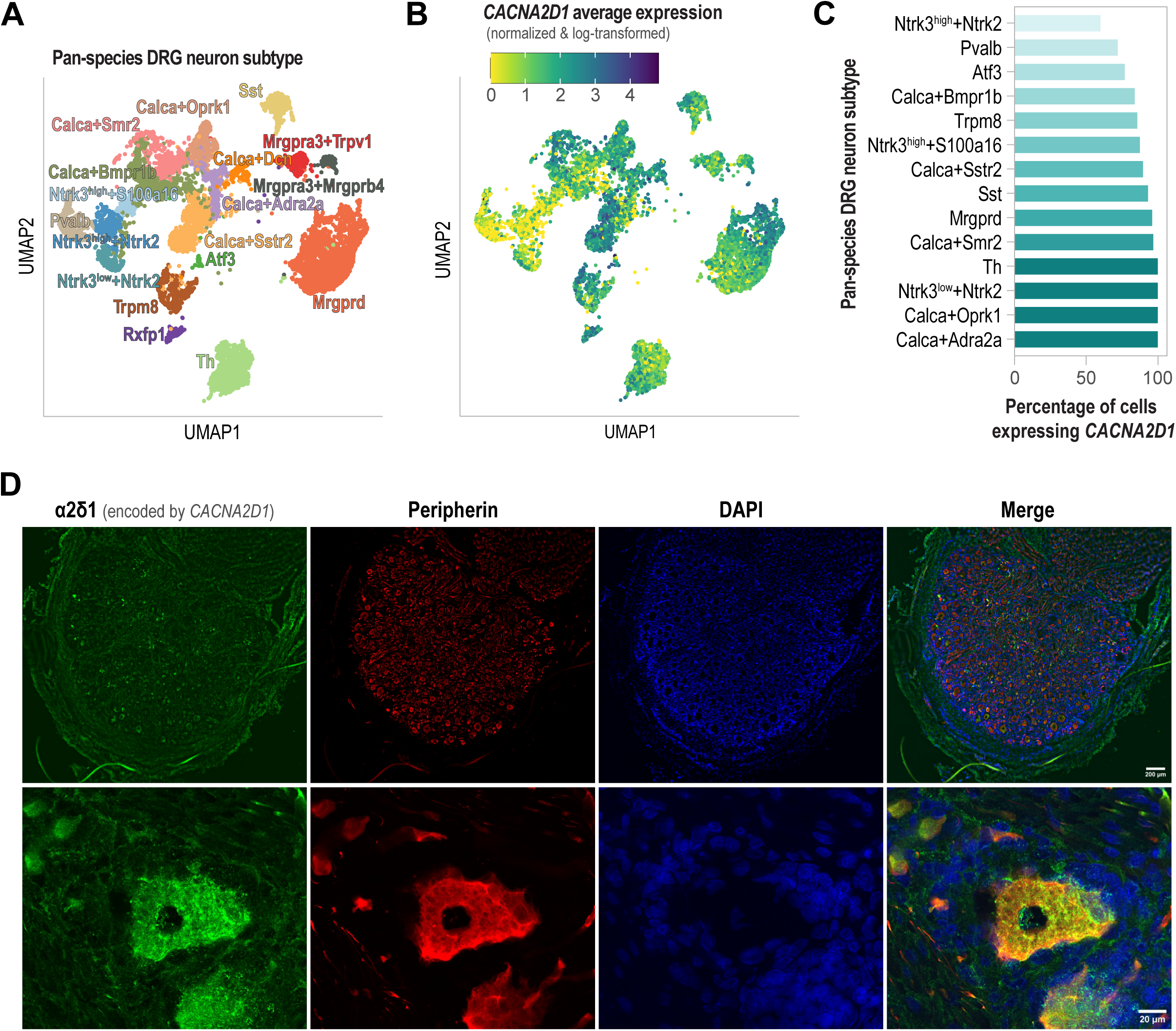
Target of GBP, α2δ1, is highly expressed in hDRG neurons. (A) UMAP plot depicting clustering of pan-species DRG-N based on marker gene expression, identifying diverse neuronal subtypes24. (B) UMAP overlay showing the expression of CACNA2D1 (encoding α2δ1) across neuronal subtypes, indicating widespread and consistent expression24. (C) Quantification of CACNA2D1 expression across neuronal subtypes24. (D) Immunohistochemical staining of hDRG tissue for α2δ1 (green), peripherin (red, nociceptor marker), and DAPI (blue, nuclear stain). Merged images confirm robust expression of α2δ1 in peripherin-positive nociceptive neurons.

To confirm the expression of α2δ1 protein in human DRG neurons, we conducted IHC on DRG tissues obtained from human donors. Peripherin, a well-established marker for nociceptors [41,42,50] was used for co-staining. IHC revealed robust α2δ1 expression in cells positive for peripherin, highlighting the receptor’s presence in nociceptive neurons (Figure 1D). Together, these results confirm that α2δ1 is abundantly expressed in human DRG neurons, spanning multiple neuronal subtypes, and is readily detectable in nociceptors. These findings reinforce α2δ1’s potential role as a key mediator of GBP’s effects on neuronal excitability and pain modulation.

### Electrophysiological properties of hDRG neurons treated with GBP

#### Effects on firing pattern prevalence, excitability, and spontaneous activity

To examine the effect of GBP on hDRG, we first analyzed the actions of *in vitro* GBP (100 µM, overnight treatment) on hDRG-N using whole-cell path-clamp electrophysiology. We chose overnight treatment as GBP actions are reported to take at least 17 hrs to develop *in vitro* and days to weeks in a clinical setting[4]. Given the limited tissue availability for recording per donor, it was only possible to perform studies using a single dose and time course. We characterized the actions of GBP on hDRG-N based on firing pattern i.e. single, multi-, rebound, or delayed firing as described previously [53]. Though the proportion of single versus multi-firing cells did not change after GBP treatment (Figure 2A), several other phenotypic properties associated with neuronal excitability changed after overnight GBP treatment in hDRG-N. Rebound firing was completely abolished in our hDRG-N data set after GBP treatment (0%, 0/21) compared to control cells (16%, 5/39), p < 0.0001 (Figure 2B). GBP treatment also reduced the prevalence of delayed firing, FSL > 100 ms, (14%, 3/21) compared to control cells (27%, 10/39), p = 0.0228 (Figure 2C).

**Figure 2:**
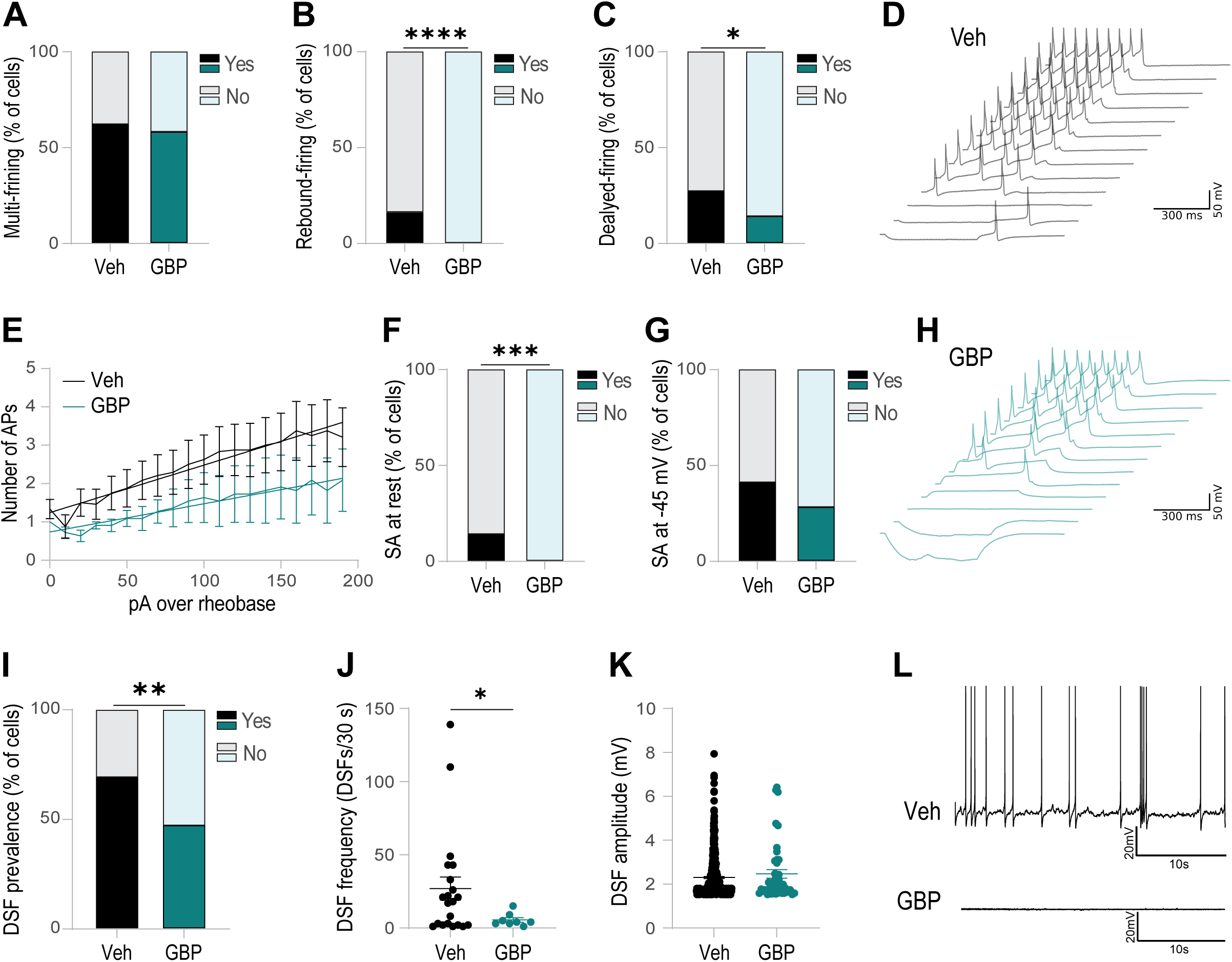
GBP reduces neuronal excitability in hDRG neurons. (A) Percentage of multi-vs. single-firing hDRG-N. (B) Percentage of cells with rebound firing. (C) Percentage of cell with delayed firing, FSL > 100 ms. (D) Example traces of a multi-firing vehicle-treated cell. (E) Frequency-current (f-I) relationship in multi-firing cells. (F) Percentage of cells with spontaneous activity (SA) at resting membrane potential. (G) Percentage of cells with SA when held at-45 mV. (H) Example traces of a multi-firing GBP-treated cell. (I) Percentage of cells with DSFs. (J) DSF frequency/30s. (K) DSF amplitudes. (L) Example traces of a vehicle-treated cell with SA and DSFs, and a GBP-treated cell with no SA and no DSFs. *p < 0.05, **p < 0.01, ***p < 0.001, ****p < 0.0001, by Fisher’s exact test for (B), (C), (F), and (I). By Kolmogorov-Smirnov for (J).

When looking at frequency-current (number of APs vs. current injection) relationships between control and GBP-treated hDRG-N, the slopes of the linear regression were not significantly different; however, the elevations of the lines were significantly different, p<0.0001, showing that multi-firing GBP-treated cells were less excitable than control cells (Figure 2E). Spontaneous activity at rest was also completely abolished in GBP-treated hDRG-N (0%, 0/21) when compared to control hDRG-N (14%, 4/39), p = 0.0001 (Figure 2F, example traces Figure 2L). Spontaneous activity when enough current was applied to the cell to hold the membrane voltage at-45 mV (also called ongoing activity) was reduced in GBP-treated cells (28%, 6/21) compared to control cells (41%, 14/39), but the difference was not statistically significant, p = 0.0531 (Figure 2G). DSF prevalence was reduced in GBP-treated hDRG-N (47%, 817) compared to control cells (69%, 22/32), p = 0.0016 (Figure 2I). DSF frequency in cells that had DSFs was also reduced in GBP-treated hDRG-N (5.50 ± 1.58 DSF/30 sec) compared to control cells (26.90 ± 7.89 DSF/30 sec), p = 0.0455 (Figure 2J, example traces Figure 2L). The DSF amplitude was not significantly different between control and GBP-treated cells (Figure 2J). Rebound firing, increased firing frequency, spontaneous activity, and increased DSF prevalence/ frequency are all associated with increased neuronal excitability. Since GBP reduced these phenotypes in hDRG-N, these data show that GBP treatment reduces neuronal excitability in hDRG-N.

#### Effects on intrinsic properties

Overnight GBP treatment had a greater effect on reducing neuronal excitability in multi-firing hDRG-N than single firing hDRG-N. In single-firing hDRG-N, the only significantly different electrophysiological property was a reduction in input resistance of GBP-treated hDRG-N (42.34 ± 13.41 MΩ) compared to control hDRG-N (148.3 ± 45.41 MΩ), p = 0.0337 (Figure 3C). In single firing hDRG-N, there was no significant change in other intrinsic properties including RMP (Figure 3B, example traces Figure 3A), rheobase (Figure 3D), normalized rheobase (Figure 3E), AP peak (Figure 3F), AP rise time (Figure 3G), or AP max rise slope (Figure 3H). In contrast, multi-firing hDRG-N had several changes in electrophysiological properties tied to neuronal excitability. When measuring RMP, GBP-treated multi-firing hDRG-N were hyperpolarized (-57.26 ± 1.514 mV) compared to control hDRG-N (-46.09 ± 2.25 mV), p = 0.0021 (Figure 3J, example traces Figure 3I). GBP-treated multi-firing hDRG-N also had an increased rheobase and normalized rheobase (371.5 ± 103.4 pA and 6.073 ± 2.268 pA/pF) compared to control hDRG-N (163.0 ± 34.5 pA and 1.784 ± 0.370 pA/pF), p = 0.0199 and p = 0.0234 respectively (Figure 3L and 3M). AP Peak was increased after GBP treatment (60.54 ± 1.26 mV) compared to control cells (52.74 ± 2.26 mV), p = 0.0179 (Figure 3N). AP rise time was significantly reduced in GBP-treated cells (1.246 ± 0.259 ms) compared to control multi-firing hDRG-N cells (1.522 ± 0.117 ms), p = 0.3010 (Figure 3O). Max rise slope was also increased in GBP-treated hDRG-N compared to control cells (163.2 ± 18.54 dVm and 116.0 ± 14.19 dVm respectively), though it was just shy of being significantly different, p = 0.0503. Input resistance was not significantly different in control or GBP-treated multi-firing cells (Figure 3K). Hyperpolarized RMP and an increased rheobase are indications of reduced neuronal excitability; therefore, by these measurements, GBP reduces the neuronal excitability in multi-firing hDRG-N but not single-firing hDRG-N. Additionally, GBP also had an effect on AP waveform properties exclusively in multi-firing cells, specifically reducing the AP rise time and AP peak.

**Figure 3:**
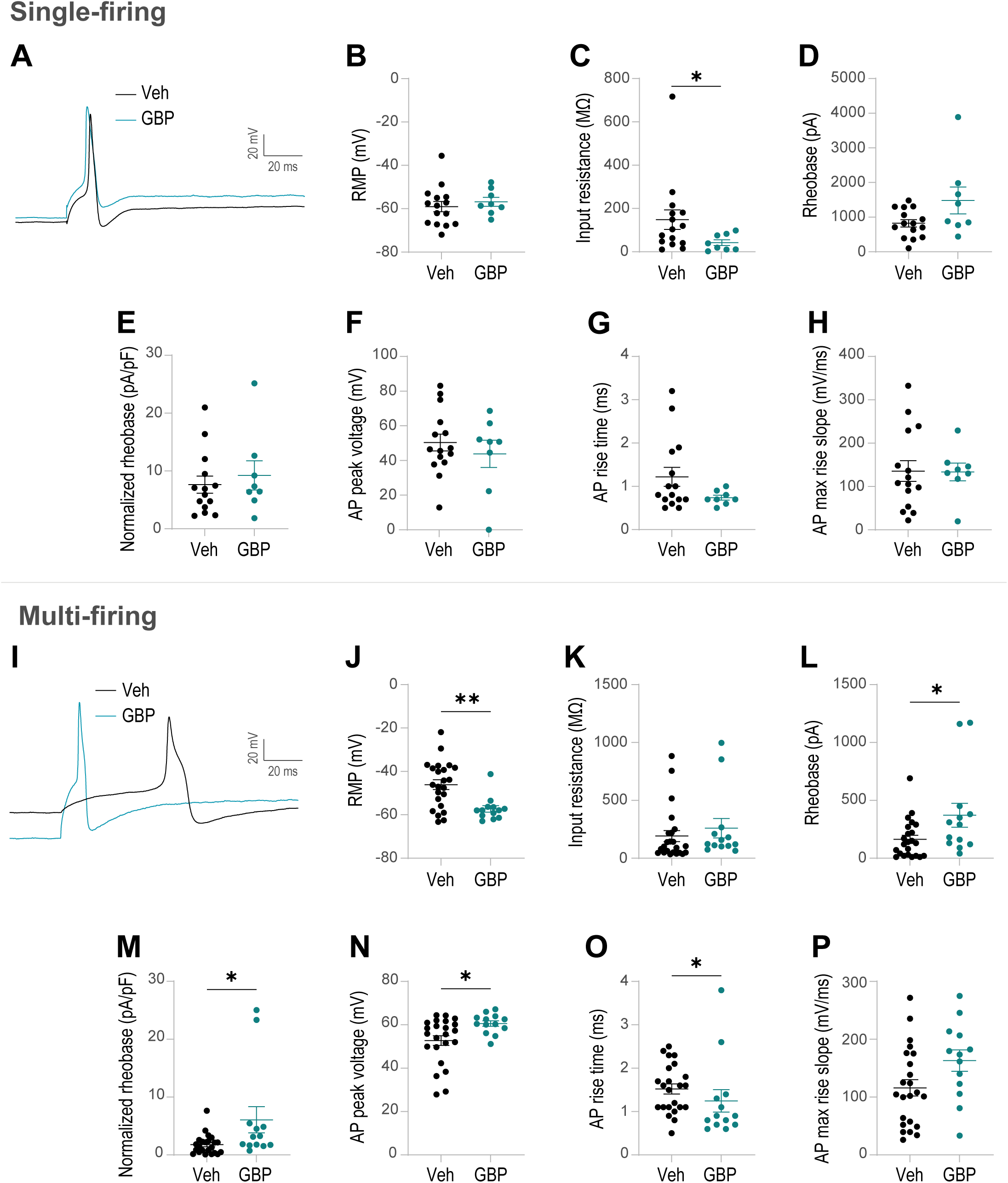
GBP reduces the excitably of multi-firing hDRG neurons but not single-firing hDRG neurons. Top half of the figure: single-firing hDRG-N. (A) Example of the rheobase trace of single-firing hDRG-N; black trace is vehicle-treated while teal trace is GBP-treated. (B) Resting membrane potential (RMP). (C) Input resistance. (D) Rheobase. (E) Normalized rheobase. (F) AP peak. (G) AP rise time. (H) AP max rise slope. Bottom half of the figure: multi-firing hDRG-N. (I) Example traces of a rheobase spike of multi-firing hDRG-N; black trace is vehicle-treated while teal trace is GBP-treated. (J) RMP. (K) Input resistance. (L) Rheobase. (M) Normalized rheobase. (N) AP peak. (O) AP rise time. (P) AP max rise slope. *p < 0.05, **p < 0.01, by Mann-Whitney.

### Donor-specific electrophysiological effects of GBP

We found that GBP treatment had a greater effect on the electrophysiological properties related to neuronal excitability on a specific donor, H22, compared to donors H16 and H17. At baseline, H22 was more excitable than H16 and H17 by several measures. H22 had a higher RMP than H16/H17,-44.95 ± 3.67 mV vs.-54.06 ± 2.08 mV, p = 0.0066 (Figure 4A). H22 also had a higher prevalence of rebound firing (3/12 hDRG-N, 25%) than H16/H17 (3/26 hDRG-N, 8%), p = 0.0020 (Figure 4E), prevalence of spontaneous activity at rest (3/12, 25% for H22, 3/26, 8% for H16/H17, p =0.0020) (Figure 4F), and prevalence of multi-firing cells (9/12, 75% for H22, 14/26, 54% for H16/H17, p = 0.0030) (Figure 4H). Taken together, these data show that hDRG-N from H22 were different from H16/H17 at baseline, demonstrating donor-specific differences in electrophysiological properties related to neuronal excitability.

**Figure 4:**
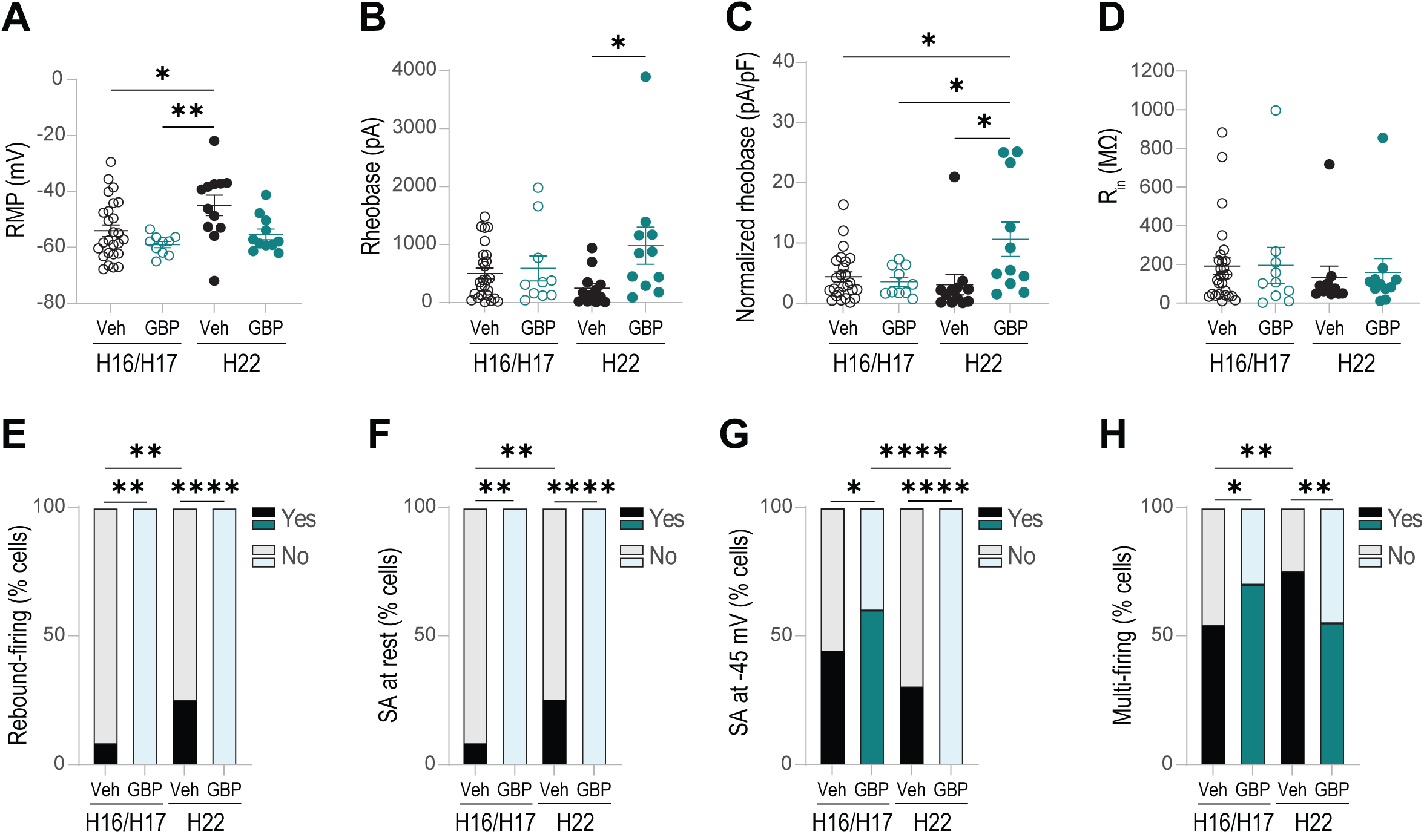
Donor specific electrophysiological effects of GBP. (A) RMP. (B) Rheobase. (C) Normalized rheobase. (D) Input resistance (Rin). (E) Percentage of cells with rebound firing. (F) Percentage of cells with SA at rest. (G) Percentage of cells with SA at-45 mV. (H) Percentage of cells with multi-firing. *p<0.05, **p<0.01, ****p<0.0001 by one-way ANOVA for (A), (B), (C), and (D). By Fisher’s exact test for (E), (F), (G), and (H).

Our analysis also showed that H22 was more responsive to GBP treatment compared to H16/H17. GBP-treated hDRG-N from H22 showed a higher rheobase (980.9 ± 319.3 pA) and normalized rheobase (10.63 ± 2.86 pA/pF) compared to control hDRG-N from H22 (rheobase of 249.2 ± 85.1 pA, normalized rheobase of 3.094 ± 1.662 pA/pF), p = 0.0357 and p = 0.0113 respectively (Figure 4B and 4C). There was no significant change in rheobase or normalized rheobase in H16/H17 after GBP treatment (Figure 4B and 4C). GBP treatment was able to reduce rebound firing and spontaneous activity in both H22 and H16/H17, though the effect was greater in H22 (Figure 4E and Figure 4F). While GBP treatment did not reduce ongoing activity in H16/H17 (Figure 4G), it completely abolished ongoing activity in H22 compared to control hDRG-N for H22 (3/10, 30% for control and 0/11, 0% for GBP-treated cells), p < 0.0001 (Figure 4G). GBP treatment was also able to reduce the prevalence of multi-firing in H22 (6/11, 54%) compared to control hDRG-N (9/12, 75%), p = 0.0047, though no reduction in multi-firing was seen in H16/H17 (Figure 4H). Though GBP treatment had a moderate effect on decreasing excitability in H16/H17 when looking at rebound firing and spontaneous activity, GBP’s effect on reducing neuronal excitability was far greater in H22; it affected more electrophysiological measurements, including increasing rheobase and normalized rheobase and decreasing rebound firing, spontaneous activity, ongoing activity, and multi-firing. We therefore termed H16/H17 as non-responders and H22 as a responder through the rest of the analysis. These data show there is a donor-specific effect of GBP treatment on hDRG-N, which may explain patient-specific effectiveness of GBP [31].

### Donor-specific hDRG transcriptomic profile associated with GBP responsiveness

Given electrophysiological differences in the baseline properties and response to GBP in H22 versus H16/H17 hDRG-N, we conducted RNA-seq on hDRG tissue from these three donors to elucidate unique transcriptomic signatures in H22 (termed the “responder” from hereon in) relative to H16/H17 (the “non-responders”). As expected, PCA analysis of the three transcriptomes revealed that the responder had a unique transcriptomic profile relative to the non-responders, the latter of which clustered together, especially along PC1 (Figure 5A). This finding supported the relevance of further inquiry into the transcriptomic differences in hDRG of the responder relative to the non-responders.

**Figure 5:**
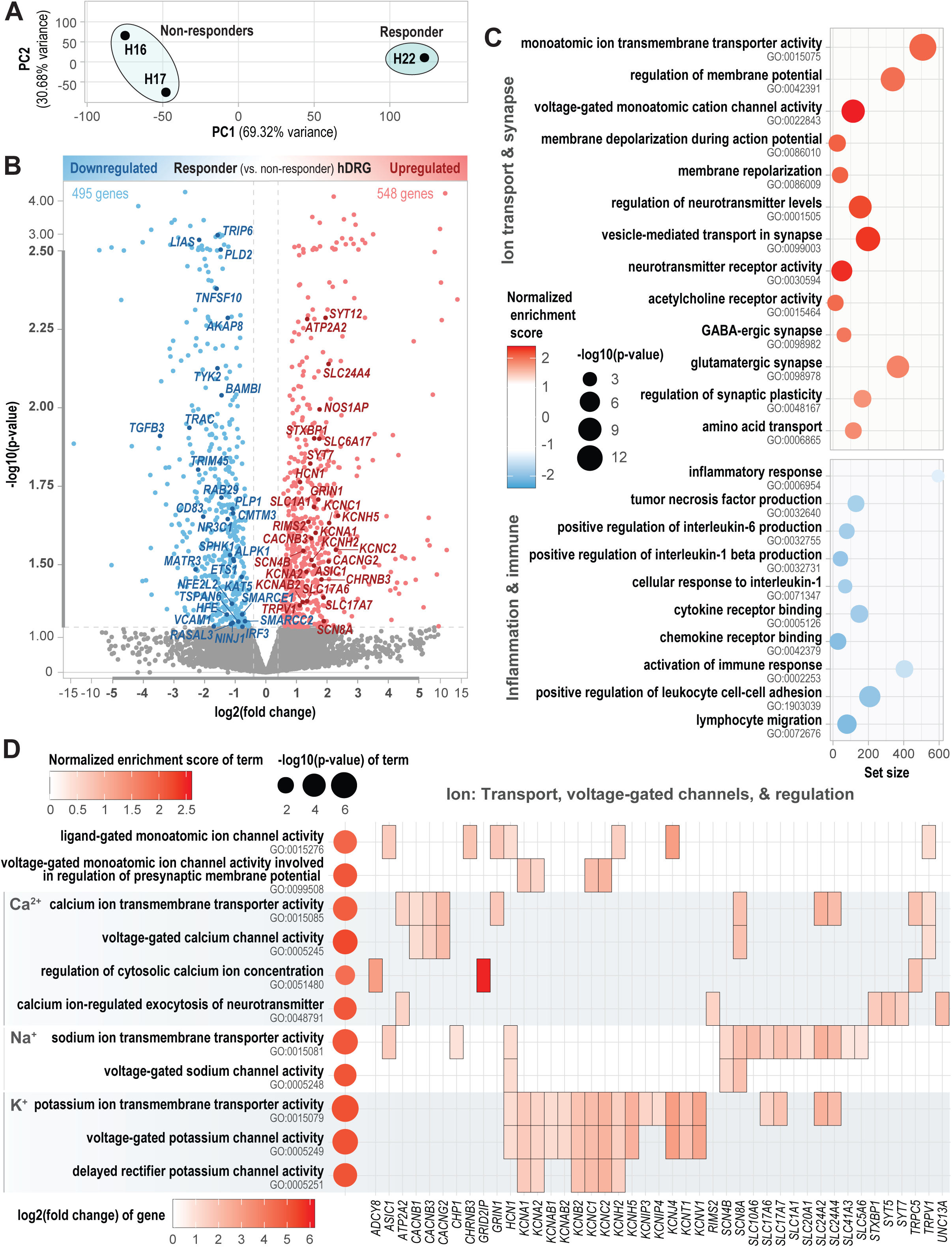
Donor-specific hDRG transcriptomic profile associated with GBP responsiveness. (A) PCA analysis of gene expression in hDRG tissue of the donor that responded electrophysiologically to GBP (H22, the “responder”) and the donors that did not respond (H16/H17, the “non-responders”). (B) Volcano plot representing the differential gene expression analysis of the responder hDRG vs. non-responder hDRGs. Significant DEGs (|log2FC| ≥ 0.4, p ≤ 0.05) in the responder vs. non-responders are colored red (upregulated) or blue (downregulated). DEGs that are core genes contributing to the enrichment of the GSEA terms shown in (C) are labeled. Labeled upregulated genes contribute to at least 3 of the positively enriched GSEA terms in (C) while downregulated genes contribute to at least 1 negatively enriched term. The thicker central portions of the axes are expanded relative to the peripheral thinner portions. (C) GSEA analysis of GO terms comparing responder vs. non-responder hDRGs. Select significant GO terms (p ≤ 0.05) relate to ion transport, synaptic transmission, inflammation, and immune response. The x-axis represents the set size (number of expressed genes in the term). Dot color represents the normalized enrichment score while dot size represents-log10(p-value). (D) GSEA analysis of GO terms related to ions (their transport, voltage-gated channels, and regulation), specifically focusing on calcium, sodium, and potassium. Dot plot on the left represents the statistics associated with the terms; dot color represents the normalized enrichment score while dot size represents-log10(p-value). The heatmap on the right represents the DEGs that constitute part of the core enrichment of the terms, with tile color indicative of log2FC of the DEGs.

Differential expression analysis of hDRG tissue from the responder versus non-responders identified 548 upregulated DEGs (log2FC ≥ 0.4, p ≤ 0.05) and 495 downregulated DEGs (log2FC ≤-0.4, p ≤ 0.05) (Figure 5B). Additionally, gene set enrichment analysis (GSEA) using Gene Ontology (GO) terms revealed transcriptome-wide differences in the expression of genes related to ion transport, synaptic transmission, inflammation, and immune response in the hDRG of the responder versus non-responders (Figure 5C). Overall, genes associated with key neuronal processes—i.e., ion transport, membrane polarization, and synaptic signaling—were upregulated whereas genes associated with neuroinflammation and immune response were downregulated (Figure 5C).

For example, the responder exhibited positive enrichment of genes involved in neuronal processes including ion transmembrane transporter activity (e.g., DEGs including *KCNV1*, *ATP2A2*, and *NIPAL3*) and regulation of membrane potential (e.g., *ABAT*, *ATP2A2*, *SLC24A4*) as well as voltage-gated cation channel activity (e.g., *KCNV1*, *KCNB2*, *KCNAB1*), membrane depolarization (e.g., *SLMAP*, *SCN4B*, *KCNH2*), and membrane repolarization (e.g., *SLC24A4*, *NOS1AP*, *KCNA1*) (Figure 5B-C). The responder also demonstrated positive enrichment of genes involved in synaptic activity, including regulation of neurotransmitter levels (e.g., *PTPRN2*, *SYT12*, *ATP2A2*), vesicle-mediated synaptic transport (e.g., *SYT12*, *ATP2A2*, *RPH3A*), neurotransmitter receptor activity (e.g., *GRIN1*, *GPR158*, *CHRNB3*), regulation of synaptic plasticity (e.g., *SYT12*, *STXBP1*, *ADCY8*), and amino acid transport (e.g., *ABAT*, *PRKG1*, *STXBP1*) (Figure 5B-C). Genes specifically associated with cholinergic (e.g., *CHRNB3*), GABAergic (e.g., *SLC6A17*, *IQSEC3*, *SST*), and glutamatergic (e.g., *PLXNA4*, *NPTX2*, *CDH8*) synapses were enriched (Figure 5B-C). Together, these gene set differences suggest a unique molecular landscape associated with ion transport, membrane potential changes relating to action potentials, and synaptic transmission, which may directly relate to the observed donor-specific electrophysiological baseline properties and GBP response (Figure 4).

Given the central importance of ion transport and voltage-gated ion channels in electrophysiological activity, we further examined gene set changes related to these features. In the responder hDRG, there was enrichment of genes related to ligand-gated ion channel activity (e.g., DEGs including *HCN1*, *GRIN1*, and *KCNJ4*) and voltage-gated ion channels involved in regulation of presynaptic membrane potential (e.g., *KCNC1*, *KCNA1*, *KCNC2*) (Figure 5D). Gene expression specifically related to the transmembrane transport and voltage-gated channels of calcium (e.g., *ATP2A2*, *SLC24A4*, *SLC24A2*, *CACNB3*, *CACNB1*, *CACNG2, TRPV1*), sodium (e.g., *SLC24A4*, *SLC24A2*, *HCN1*, *CHP1*, *SCN4B*, *SCN8A*), and potassium (e.g., *KCNV1*, *SLC24A4*, *KCNB2*, *KCNIP3*, *KCNAB1*, *HCN1*) was strongly upregulated (Figure 5D). Finally, there was also enrichment of genes involved in regulating cytosolic calcium concentration (e.g., *ADCY8*, *TRPC5*, *GRID2IP*) and calcium-regulated neurotransmitter exocytosis (e.g., *ATP2A2*, *STXBP1*, *SYT7*) (Figure 5D). Ultimately, over 40 upregulated DEGs contributed to these ion-related terms (Figure 5D). These results indicate widespread upregulation of genes related to ion channels—especially of calcium, sodium, and potassium—and related their functions in the hDRG of the responder.

On the other hand, the responder hDRG gene set was relatively deficient in the expression of genes associated with inflammatory and immune responses (Figure 5C). For example, the responder exhibited negative enrichment of genes associated with inflammatory mediator production, including tumor necrosis factor (TNF) (e.g., DEGs including *AKAP8*), interleukin-6, and interleukin-1 beta (Figure 5B-C). Additionally, genes involved in cytokine (e.g., *TRIP6*, *TNFSF10*, *TYK2*) and chemokine receptor binding, immune response activation (e.g., *PLD2*, *TRAC*, *RAB29*), leukocyte cell-cell adhesion (e.g., *TYK2*, *CD83*, *ETS1*), and lymphocyte migration were collectively downregulated (Figure 5B-C). These findings indicate that the responder hDRG exhibited dampened gene expression related to inflammation and immune system activity, which may translate to functional differences in these responses.

### Transcriptomic properties of hDRG treated with GBP

Given the baseline transcriptomic differences between the responder and non-responders, we hypothesized that GBP-induced electrophysiological changes in the responder hDRG-N may be coupled with and potentially driven by transcriptomic alterations. Therefore, we performed RNA-seq on cultured hDRG from the responder to characterize the transcriptomic changes induced by GBP that accompany electrophysiological response. Overnight GBP treatment induced the upregulation (log2FC ≥ 0.4, p ≤ 0.05) of 144 genes and downregulation (log2FC ≤-0.4, p ≤ 0.05) of 47 genes (Figure 6A). GSEA revealed transcriptome-wide alterations in the expression of genes associated with the same overarching processes that were altered in the responder versus non-responders: ion transport, synaptic transmission, inflammation, and immune response (Figure 6B).

**Figure 6:**
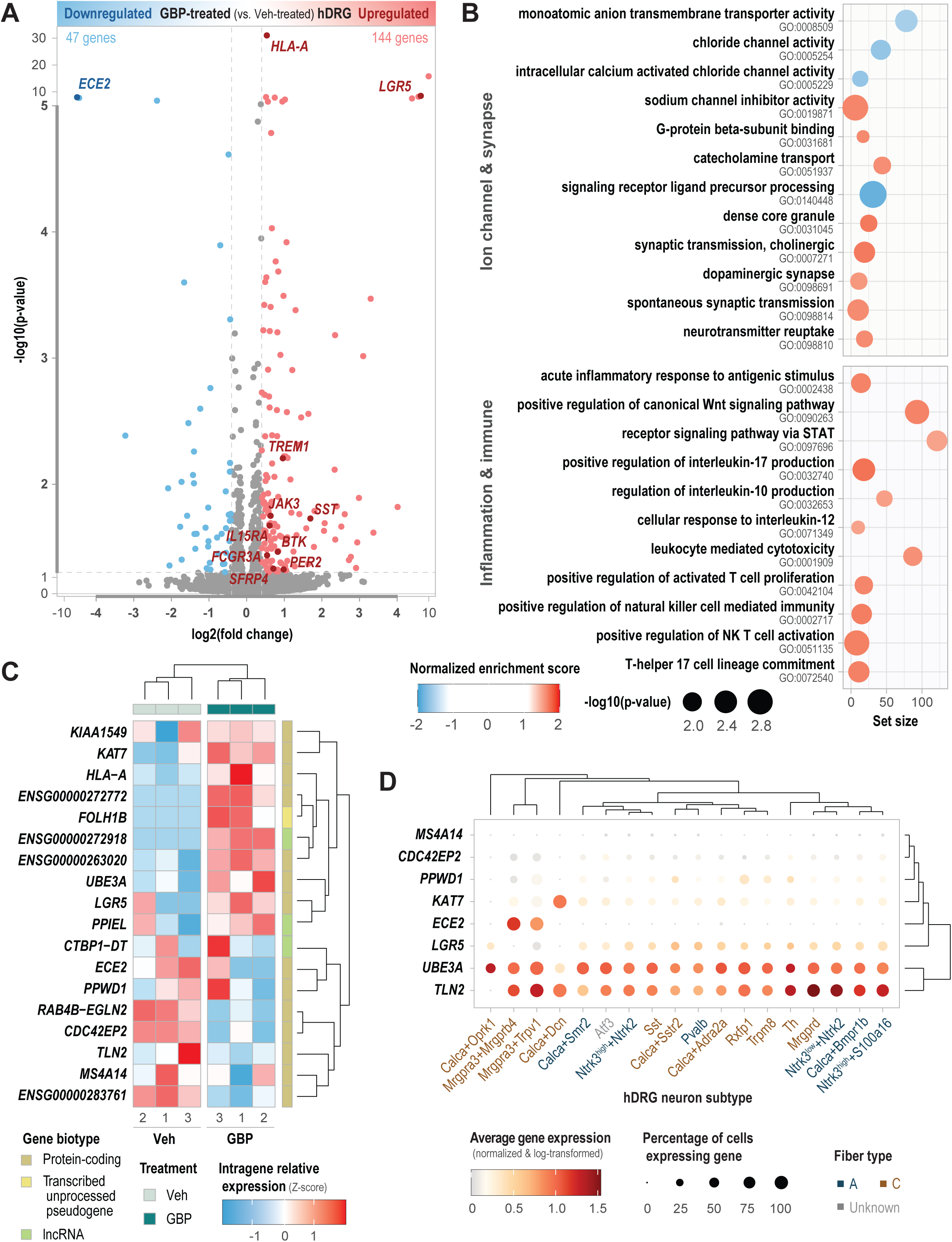
Transcriptomic properties of hDRG treated with GBP. (A) Volcano plot representing the differential gene expression analysis of the responder hDRG treated *in vitro* with GBP vs. vehicle (Veh). Significant DEGs (|log2FC| ≥ 0.4, p ≤ 0.05) in GBP-treated vs. vehicle-treated hDRG are colored red (upregulated) or blue (downregulated). DEGs that are core genes contributing to the enrichment of at least one GSEA term shown in (B) are labeled. The thicker central portions of the axes are expanded relative to the peripheral thinner portions. (B) GSEA analysis of GO terms comparing GBP-treated vs. vehicle-treated hDRG culture. Select significant GO terms (p ≤ 0.05) relate to ion transport, synaptic transmission, inflammation, and immune response. The x-axis represents the set size (number of expressed genes in the term). Dot color represents the normalized enrichment score while dot size represents-log10(p-value). (C) Heatmap depicting the relative expression of strong DEGs (|log2FC| ≥ 0.4, FDR ≤ 0.1) among the 3 vehicle-treated and 3 GBP-treated hDRG samples. Color represents relative expression within each gene (Z-score). Gene biotype and treatment are annotated by color, and hierarchical clustering of samples and genes are shown in their respective dendrograms. (D) Expression of strong DEGs (|log2FC| ≥ 0.4, FDR ≤ 0.1) in hDRG-N subtypes as quantified and characterized by previous studies24. Dot color represents average gene expression (normalized and log-transformed counts) while dot size represents the percentage of cells expressing the gene. Text color of hDRG-N subtype labels indicate their fiber type.

Compared to control hDRG cultures, GBP-treated cultures were deficient in the expression of genes involved in anion transmembrane transport, including the activity of chloride channels and specifically intracellular calcium-activated chloride channels (Figure 6A-B). Meanwhile, GBP treatment resulted in the positive enrichment of genes involved in inhibiting cation flux, including sodium channel inhibitor activity, G-protein beta-subunit binding, and catecholamine transport. GBP also tended to enhance the expression of genes involved in aspects of synaptic signaling, such as ligand precursor processing (e.g., DEGs including *ECE2*), dense core granules (e.g., *SST*), spontaneous synaptic transmission, and neurotransmitter reuptake (e.g., *PER2*) (Figure 6A-B). This particularly included genes found in cholinergic and dopaminergic synapses (Figure 6A-B). Together these gene set differences suggest that GBP may have suppressive actions on ion transport but enhance certain synaptic activities.

GBP treatment also led to the upregulation of genes involved in inflammatory and immune processes (Figure 6B). The hDRG culture treated with GBP exhibited positive enrichment of genes involved in acute inflammatory response to antigenic stimulus (e.g., DEGs such as *BTK*, *FCGR3A*), inflammation-related signaling pathways—canonical Wnt (e.g., *LGR5*, *SFRP4*) and STAT (e.g., *JAK3*, *IL15RA*)—and interleukin production (e.g., *JAK3*, *BTK*) and response. GBP also induced the upregulation of genes associated with the activation and function of several immune cell types, such as leukocytes (e.g., *HLA-A*, *TREM1*, *FCGR3A*), T cells, natural killer cells, CD8+ T cells, and T-helper 17 cells (Figure 6A-B).

Strong DEGs (|log2FC| ≥ 0.4, FDR ≤ 0.1) contributing to enriched terms included *HLA-A* (log2FC = 0.54, p = 9.27e-32, FDR = 1.50e-27; leukocyte mediated cytotoxicity), *LGR5* (log2FC = 8.49, p = 3.24e-9, FDR = 1.74e-5; positive regulation of canonical Wnt signaling pathway), and *ECE2* (log2FC =-7.59, p = 1.06e- 8, FDR = 2.85e-5; signaling receptor ligand precursor processing) (Figure 6B-C). Other strongly upregulated genes (log2FC ≥ 0.4, FDR ≤ 0.1)—all protein-coding unless otherwise indicated—included *FOLH1B* (transcribed unprocessed pseudogene), *TLN2*, *PPWD1*, *KIAA1549*, *CTBP1-DT* (lncRNA), *UBE3A*, *KAT7*, and *PPIEL*. Meanwhile, other strongly downregulated genes (log2FC ≤-0.4, FDR ≤ 0.1) included *MS4A14*, *RAB4B-EGLN2*, and *CDC42EP2* (Figure 6C). Based on the expression of strong DEGs, the VEH and GBP hierarchically clustered together, highlighting the defining nature of these DEGs in the response to GBP (Figure 6C).

To better grasp the physiological relevance of these strong DEGs in hDRG, we examined their expression in different hDRG-N cell types using a publicly available reference combining data from three single-cell hDRG studies [7] (Figure 6D). By both average gene expression and percentage of cells expressing the gene, *TLN2* and *UBE3A* exhibit the most robust expression across neuron types in hDRG and hierarchically cluster closely together (Figure 6D). *LGR5* is also widely expressed throughout hDRG-N subtypes though to a lesser degree (Figure 6D). Meanwhile, *ECE2* is highly expressed in *MRGPRA3*- defined cell types (Mrgpra3+Mrgprb4 and Mrgpra3+Trpv1) while *KAT7* is highly expressed in Calca+Dcn neurons (Figure 6D). Ultimately, given the collectively wide expression of these strong DEGs across hDRG-N subtypes, GBP has the potential to affect a variety of hDRG-N perturbed in neuropathic pain conditions.

### Cross-transcriptomic analysis comparing GBP-responsive and GBP-treatment hDRG signatures

At baseline, the responder hDRG-N had a more excitable electrophysiological phenotype than the non-responder hDRG-N by several measures—increased RMP, increased rebound firing, and increased proportion of multi-firing cells—which was subsequently rescued by GBP treatment (Figure 4). Given also the transcriptome-wide alterations in gene expression related to similar neuronal and immunoinflammatory processes in both RNA-seq datasets, we next cross-analyzed the baseline responder-specific transcriptomic profile and the GBP-treatment profile from the responder hDRG. This enabled us to understand which processes and genes were perturbed in the responder relative to the non-responder hDRG but rescued by GBP treatment in the hDRG culture. This pattern of gene expression correlates with the enhanced baseline excitability of the responder hDRG-N that was rescued by GBP treatment.

We observed that positive enrichment of gene expression associated with certain processes related to ion transport and synaptic signaling in the responder hDRG at baseline was subsequently downregulated by GBP treatment in the responder hDRG culture (Figure 7A). Ion transport-related rescues included the expression of genes involved in the behavioral response to chemical pain, anion channel activity, and chloride transmembrane transport (Figure 7A). Select synapse-related genes also exhibited the same “up-down” rescue pattern, including genes encoding presynaptic proteins and genes involved in D-amino acid, L-glutamate, and L-aspartate transport (Figure 7A).

**Figure 7:**
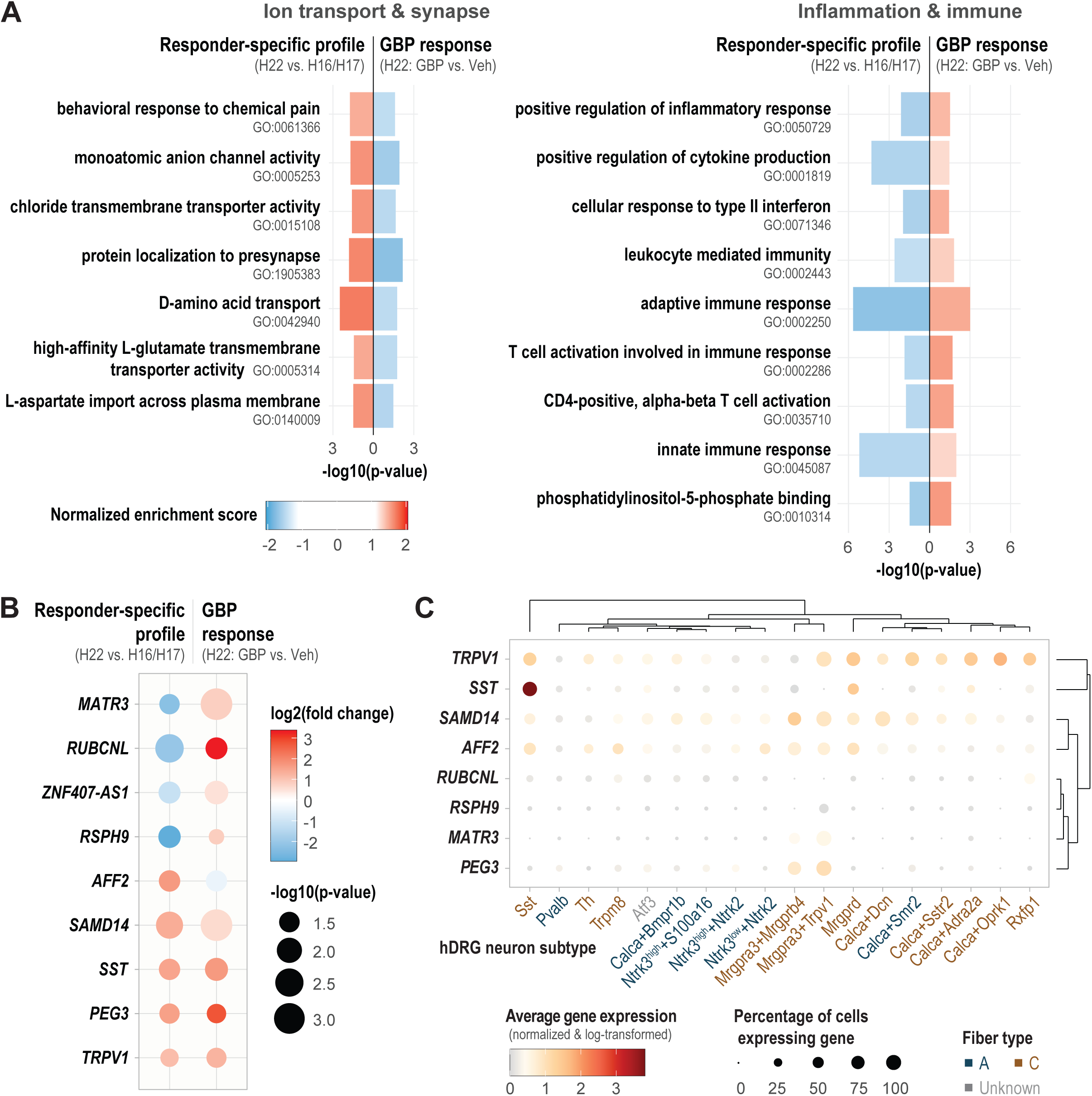
Cross-transcriptomic analysis comparing GBP-responsive and GBP-treatment signatures. (A) GO terms enriched by GSEA analysis in opposite directions in the responder vs. non-responder hDRG dataset and the GBP-treated vs. vehicle-treated hDRG dataset. Select significant GO terms (p ≤ 0.05) relate to ion transport, synaptic transmission, inflammation, and immune response. The x-axis represents-log10(p-value) bidirectionally, and the bar color represents the normalized enrichment score. (B) Common DEGs (|log2FC| ≥ 0.4, p ≤ 0.05) among the responder vs. non-responder hDRG dataset and the GBP-treated vs. vehicle-treated hDRG dataset. Dot color represents log2(fold change) while dot size represents-log10(p-value). (C) Expression of these common DEGs (|log2FC| ≥ 0.4, p ≤ 0.05) in hDRG- N subtypes as quantified and characterized by previous studies24. Dot color represents average gene expression (normalized and log-transformed counts) while dot size represents the percentage of cells expressing the gene. Text color of hDRG-N subtype labels indicate their fiber type.

Meanwhile, genes set-wide expression related to processes involved in inflammation and immune response demonstrated a rescue in the opposite direction: deficient in the responder hDRG but upregulated by GBP treatment (Figure 7A). This included the overall expression of genes involved in positively regulating inflammatory response and cytokine production as well as genes involved in type II interferon response and leukocyte-mediated immunity (Figure 7A). Additionally, transcriptome-wide downregulation of genes involved in the adaptive and innate immune responses – T helper cells and phosphatidylinositol-5-phosphate binding, respectively – were rescued by GBP treatment (Figure 7A). Together, these results suggest a GBP-induced rescue of select neuronal processes via reversing downregulated expression of genes associated with immune responses.

Several genes exhibited a rescue pattern as significant DEGs (|log2FC| ≥ 0.4, p ≤ 0.05) in both datasets. *MATR3*, *RUBCNL*, *ZNF407-AS1*, and *RSPH9* were deficient in the responder relative to the non-responder hDRG then upregulated by GBP treatment in the responder hDRG, following the rescue trend of the immunoinflammatory terms (Figure 7B). Notably, *MATR3* was a core gene contributing to the opposing enrichment of the innate immune response term while *RUBCNL* similarly contributed to the enrichment of phosphatidylinositol-5-binding (Figure 7A-B). Conversely, *AFF2* was more highly expressed in the responder hDRG then downregulated by GBP treatment (Figure 7B). Additionally, *SAMD14*, *SST*, *PEG3*, and *TRPV1* were more abundantly expressed in the responder than non-responder hDRG then further upregulated by GBP treatment (Figure 7B).

Eight of the nine DEGs that appeared in both the responder and GBP-treatment profile were detected in the single-cell hDRG-N public dataset previously described [7]. All eight genes tend to be expressed to a greater degree—by average expression and percentage of cells expressing the gene—in C-fiber hDRG-N subtypes than A-fiber subtypes, with Mrgpra3+Trpv1, Mrgprd, and Sst subtypes as key types expressing these genes (Figure 7C). *TRPV1* and *SST* hierarchically cluster closely together based on the similarity of their expression pattern across hDRG-N subtypes (Figure 7C). Notably, they are also key genes used to define subtypes: *SST* is very strongly expressed in Sst hDRG-N while *TRPV1* is highly expressed in the Mrgpra3+Trpv1 hDRG-N as well as several others, such as Calca+Oprk1, Calca+Adra2a, and Mgrd1 (Figure 7C). *SAMD14* and *AFF2* also clustered together and were generally widely expressed across cell types (Figure 7C). Meanwhile, *MATR3* and *PEG3* were predominantly expressed by *MRGPRA3*-defined cell types (Mrgpra3+Mrgprb4 and Mrgpra3+Trpv1) (Figure 7C). Through the upregulation of these genes (except *AFF2*), GBP may target the expression of already perturbed genes in several C-fiber subtypes of responder hDRG-N.

## Discussion

Our findings indicate that the actions of GBP are donor-specific. Despite GBP’s widespread use in treating neuropathic pain, clinical observations show limited effectiveness, with a number needed to treat (NNT) of 6-8, and adverse effects reported in many patients [6,17,31]. hDRG-N collected from H22 were in general more excitable when compared to our previously published data on hDRG-N collected from 10 other donors when comparing multiple measures of neuronal excitability such as a higher RMP, lower rheobase, higher prevalence of rebound firing, and higher prevalence of spontaneous activity in H22 [53,54]. This suggests that clinical history of H22 may explain increased neuronal excitability to confer increased responsiveness of H22 hDRG-N to *in vitro* GBP treatment. The exact reasons for the more pronounced actions of GBP on excitability of hDRG-N from H22 compared to H16 and H17 are unclear.

One potential explanation includes differences in ion channels such as the sodium “leak” channel NALCN, implicated in setting the RMP and in pain signaling [51]. The cause of death may also contribute; H22 sustained traumatic head injuries due to a motor vehicle accident, whereas the others did not. Since this is a traumatic cause of death that may involve significant tissue injury, it is possible that this may explain the specific actions of GBP. It has previously been shown that α2δ1 is upregulated in DRG and spinal cord following nerve injury [26,27], however we did not observe a significant upregulation of α2δ1 mRNA in hDRG from H22 compared to H16 and H17. Further study will require examination of α2δ1 protein levels in DRG from these donors and possibly also cell-type specific levels of α2δ1 across donors.

It is well known that the main mechanism of action of GBP in sensory neurons is by binding to the α2δ1 subunit to reduce trafficking and expression of voltage-gated calcium channels causing a reduction in neurotransmitter release and excitability [13,20]. α2δ1 knock-out mice show reduced neuronal excitability in multi-firing DRG neurons by reducing the frequency of AP firing [29]. This study also showed a decrease in the first spike latency of DRG neurons [29], which correlates with our data as we saw a reduction in hDRG-N with delayed firing, FSL > 100ms, following GBP treatment. Previous work also showed that DRG neurons from α2δ1 KO mice had shorter AP durations [29]. Our study showed a decrease specifically in AP rise time after GBP treatment in hDRG-N, though we did not see any significant changes in AP duration (data not shown). The differences between the mouse knock-out study and our study could be related to species differences, compensatory effects of α2δ1 knockout, or off-target effects of GBP treatment. The α2δ1 knock-out study also did not look at electrophysiological differences between multi-and single-firing DRG neurons. Our results suggest that the effects of GBP are more pronounced in multi-firing DRG neurons, suggesting the role of α2δ1 on neuronal excitability may be different in different neuronal subtypes. Indeed, single cell RNA-seq data from mouse DRG neurons has shown that α2δ1 expression is higher in multi-firing neurons with longer FSL such as nociceptors and C-fiber low-threshold mechanoreceptors (C-LTMRs) [52].

Increased DSF frequency in hDRG-N is also associated with increased levels of spontaneous activity and neuropathic pain [33,34,45]. Our data shows that GBP treatment reduced both spontaneous activity, DSF prevalence, and DSF frequency in hDRG-N. Previous work shows that DSFs are dependent on extracellular calcium and sodium [45]. Other data shows that patients with a history of pain have a higher prevalence of DSFs [34]. Given these publications and our data showing GBP-induced reduction in DSF prevalence and frequency, it is likely that this is an electrophysiological mechanism of GBP’s reduction in neuropathic pain.

To investigate the molecular mechanisms underlying donor-specific variability in GBP responsiveness, we performed RNA sequencing on DRG tissue from donor H22 (the “responder”) and donors H16/H17 (“non-responders”). This analysis identified several DEGs and enriched pathways in H22 that provide key insights into the mechanisms of GBP action. Enrichment analysis revealed that DEGs were associated with ion channel function, synaptic signaling, inflammation, and immune response—all pathways previously implicated in the pathophysiology of neuropathic pain. These findings support the hypothesis that GBP’s effects on neuronal excitability may depend on its ability to modulate these biological processes.

At baseline, nociceptive DRG neurons from H22 exhibited higher activity compared to H16/H17, suggesting that H22 may have had an underlying sensitization or priming of nociceptive pathways. Although neuropathic pain was not reported in H22’s medical records, this baseline hyperexcitability aligns with hallmarks of neuropathic pain, such as heightened ion channel activity and synaptic dysregulation [2]. GBP treatment appeared to counteract these aberrant processes, rescuing gene expression patterns associated with ion channels, synaptic signaling, and inflammation.

Key DEGs included TRPV1, a well-established marker of nociceptors [7,30]. TRPV1 plays a critical role in nociceptive signaling and pain perception. Interestingly, TRPV1 was upregulated in both untreated and GBP-treated DRG from H22, raising the possibility that TRPV1 facilitates GBP’s effects. Supporting this hypothesis, studies suggest that TRPV1 may serve as a conduit for GBP entry into nociceptors, enhancing its antinociceptive efficacy [8]. Future studies should validate whether TRPV1 directly mediates GBP uptake and its subsequent effects on nociceptors.

Another DEG of interest is somatostatin (SST), a neuropeptide involved in the modulation of pain signaling [7,11,25,52]. SST acts as an inhibitory signal in the DRG, potentially reducing pain transmission by suppressing sensory neuron activity [11,18,21,25,40]. The observed increase in SST expression after GBP treatment suggests that GBP may synergize with endogenous inhibitory pathways to attenuate nociceptive signaling. A plausible mechanism is that GBP’s primary action of reducing calcium channel activity and suppressing neurotransmitter release may establish a conducive environment for SST to enhance its inhibitory effects on nociceptive signaling. While direct evidence supporting this hypothesis is currently lacking, it presents a promising avenue for future research leveraging possible synergistic effects of GBP and SST to achieve more effective pain relief.

The unique responsiveness of H22 to GBP may also be influenced by donor-specific factors such as drug exposure. H22’s history of amphetamine exposure may have primed their DRG neurons through mechanisms such as neuroinflammation [9,43] and altered ion channel activity including TRP channels [14,22]. Conversely, the diminished GBP responsiveness in H16/H17 could be partially attributed to their opioid exposure, which is known to affect DRG neuronal excitability, although differential effects are observed depending on history of injury, dose, and duration of exposure [1,12]. Gabapentinoids appear to potentiate actions of opioid clinically and recent work has shown that this mechanism depends on gabapentinoid reduction of calcium influx through L-type voltage-gated calcium channels [15]. Given the differential expression of genes involved in the transport of calcium, sodium, and potassium in H22 vs. H16/H17, including their voltage-gated channels, a variety of mechanisms may influence electrophysiological responsiveness to GBP.

The observed donor-specific enrichment of ion channel and inflammation-related pathways underscores the importance of these mechanisms in modulating GBP responsiveness. For instance, alterations in calcium channel subunits and sodium channels, which are critical for setting neuronal excitability thresholds could underlie the differences in GBP efficacy between donors. For example, the relatively elevated expression of other voltage-gated calcium channel subunits (*CACNB1*, *CACNB3*, and *CACNG2*) in the responder may have influenced calcium flux and primed the responder for their potent response to GBP, which likely involved at least in part GBP’s targeting of voltage-gated calcium channels.

This study highlights the complexity of GBP’s actions on human sensory neurons and underscores the importance of individual variability in determining therapeutic outcomes. While our findings provide valuable insights, they also raise several questions that warrant further investigation. Expanding the sample size to include a broader and more diverse cohort of donors will be critical for identifying patterns of GBP responsiveness across populations. Additionally, employing single-cell RNA sequencing would allow for a more granular understanding of how GBP affects specific neuronal subtypes, such as nociceptors, mechanoreceptors, and C-LTMRs. Longitudinal studies examining the sustained impact of GBP treatment on DRG gene expression and neuronal excitability could provide insights into its long-term efficacy and mechanisms of tolerance. Moreover, integrating proteomic and metabolomic analyses with transcriptomic data could uncover additional molecular targets and pathways that modulate GBP’s effects.

In conclusion, this study provides a comprehensive electrophysiological and transcriptomic profile of a GBP-responsive donor and elucidates key mechanisms underlying GBP’s modulation of human DRG neurons. These findings enhance our understanding of GBP’s mechanistic actions on peripheral pain-sensing neurons and offer valuable insights for optimizing its clinical use in managing neuropathic pain. Future research focusing on donor-specific factors, single-cell analyses, and combinatory therapies holds promise for improving the efficacy and precision of GBP-based treatments for neuropathic pain.

## Acknowledgements

We gratefully acknowledge support from the UNM Rainforest Innovations 2024 GAP funding (SRAA) and the Research Endowment Fund of the Department of Anesthesiology and Critical Care Medicine, UNM HSC. We also extend our deepest gratitude to the organ donors and their families, whose generous contributions made this study possible.

## Conflict of Interest Statement

The authors declare no conflicts of interest.

## Data Sharing Statement

Electrophysiological, transcriptomic, and code data generated during this study will be made available to qualified researchers upon request. Data access will require a data transfer agreement to ensure proper usage and attribution. We are in the process of submitting the transcriptomic data to the Sequence Read Archive (SRA) and the analysis code to GitHub. These repositories will provide open access to the data and tools used in this study, supporting reproducibility and transparency in research.

## Notes

### Competing Interest Statement

The authors have declared no competing interest.

